# scAURA: Alignment- and Uniformity-based Graph Debiased Contrastive Representation Architecture for Self-Supervised Clustering of Single-Cell Transcriptomics

**DOI:** 10.64898/2026.01.25.701579

**Authors:** Jubair Ibn Malik Rifat, Sarthak Engala, Serdar Bozdag

## Abstract

Single-cell RNA sequencing (scRNA-seq) allows transcriptomic profiling at single-cell resolution, providing valuable insights into cellular diversity across tissues, developmental stages, and diseases. However, accurately identifying cell types remains challenging due to the high dimensionality, sparsity, and noise inherent in scRNA-seq data. To address these challenges in cell type identification in scRNA-seq data, we introduce scAURA (single <u>c</u>ell <u>A</u>lignment- and <u>U</u>niformity-based Graph Debiased Contrastive <u>R</u>epresentation <u>A</u>rchitecture), a unified framework that integrates graph debiased contrastive learning with self-supervised clustering. We evaluated scAURA on 18 real single-cell datasets collected from six sequencing platforms spanning diverse tissue and cell types in human and mouse. scAURA outperformed all state-of-the-art (SOTA) methods in nine and eight datasets in Adjusted Rand Index (ARI) and Normalized Mutual Information (NMI), respectively. On average, scAURA obtained average ranks of 2.28 (ARI) and 2.39 (NMI) across all 13 SOTA methods, demonstrating its consistent superiority across datasets. scAURA also exhibited strong robustness to dropout noise by maintaining stable clustering performance even under increasing sparsity levels. Furthermore, in an external single-cell Alzheimer’s disease dataset, scAURA accurately clustered different cell types, identified novel cell type–specific marker genes, and inferred their potential transcriptional regulators. The source code and datasets are available at https://github.com/bozdaglab/scAURA.

## 1 Introduction

Single-cell RNA sequencing (scRNA-seq) enables transcriptomic profiling at the resolution of individual cells, providing unprecedented insights into cellular heterogeneity across tissues, developmental stages, and disease states (Cha and Lee, 2020; Potter, 2018). However, accurately clustering cell types remains a fundamental challenge due to the high dimensionality, sparsity, and inherent noise of scRNA-seq data (Kiselev et al., 2019; Zhang et al., 2023). Technical artifacts such as dropout events further obscure true biological variation and complicate the identification of distinct cellular populations (Tran et al., 2020; Qiu, 2020). Conventional clustering algorithms such as *k* -means and hierarchical clustering often perform poorly under these conditions, as they are not designed to handle zero-inflated count data or the complex nonlinear structures of single-cell transcriptomes.

To address these challenges, several specialized pipelines such as Seurat (Butler et al., 2018), SHARP (Wan et al., 2020), SC3 (Kiselev et al., 2017), SCANPY (Wolf et al., 2018), and SINCERA (Guo et al., 2015) have been widely adopted in single-cell analysis. Seurat integrates dimensionality reduction, graph-based clustering, and visualization into a unified framework, while SHARP enables rapid clustering of large-scale datasets using random projection ensembles for efficient dimensionality reduction. SC3 employs consensus clustering with gene filtering, principal component analysis (PCA), and Laplacian transformation. SCANPY offers scalable analysis for large single-cell datasets by utilizing shared nearest neighbor (SNN) modular optimization and Leiden for clustering. SINCERA applies z-score normalization followed by hierarchical clustering for cell-type identification.

Although these methods have advanced single-cell analysis, they often fail to fully capture the intrinsic data distribution and true biological variability present in scRNA-seq data. Autoencoder-based frameworks such as the Deep Count Autoencoder (DCA) (Eraslan et al., 2019) model the high variability and dropout inherent in scRNA-seq data through a zero-inflated negative binomial (ZINB) loss. The Deep Embedded Clustering (DEC) (Xie et al., 2016) algorithm was originally developed for image and text data to integrate representation learning and clustering through a Kullback–Leibler divergence (KLD)–based target refinement. scDeepCluster (Lei et al., 2023) extends DEC with a ZINB autoencoder for scRNA-seq data and jointly optimizes reconstruction and clustering objectives. Similarly, scziDesk (Chen et al., 2020) replaces DEC’s KLD-based objective with a soft self-training *k* -means strategy that improves clustering stability under sparsity and noise. Other models such as DESC (Li et al., 2020) and scCAEs (Hu et al., 2022) further refine latent representations or capture nonlinear gene–gene relationships, while the probabilistic framework single-cell Variational Inference (scVI) (Lopez et al., 2018) employs variational autoencoders to model biological variability and batch effects. However, these methods remain heavily dependent on reconstruction loss, making their performance sensitive to the initial quality of the data, and they often overlook explicit modeling of cell–cell relationships, which is crucial for capturing complex biological structures.

To explicitly model cell–cell relationships, several graph-based methods have been proposed. Single-cell Graph Neural Network (scGNN) (Wang et al., 2021), Single-cell Graph Autoencoder (scGAE) (Luo et al., 2021), GraphSCC (Zeng et al., 2020), and scMGCA (Yu et al., 2023) leverage graph convolutional or autoencoder architectures to propagate information through cell similarity networks. However, their performance strongly depends on the quality of the constructed graph, which may include spurious edges introduced by technical noise or batch effects. Additionally, these methods typically construct cell graphs using a fixed *k* value for the *k* -nearest neighbors (kNN) approach, which can fail to capture rare or small cell populations due to uniform neighborhood sizes. Graph contrastive learning directly addresses the noisy edge issue by learning structure-invariant and noise-robust representations through graph augmentations, enabling the model to reduce the influence of erroneous edges and better preserve true biological relationships.

Motivated by these insights, we propose scAURA, a unified framework that integrates graph debiased contrastive learning with self-supervised clustering for scRNA-seq analysis. scAURA learns latent representations that are robust to noisy or biased graph construction while iteratively refining cluster assignments through self-training. Specifically, our contributions are:

1. We utilized an adaptive *k* -nearest neighbor (kNN) strategy that dynamically adjusts neighborhood size to better capture rare or small cell-type clusters.
2. We reduced the influence of spurious edges by combining SNN-based edge weighting with a debiased contrastive learning objective.
3. We designed a modified contrastive loss that incorporates alignment and uniformity to ensure that embeddings of similar cells are pulled closer while maintaining a well-dispersed latent distribution.
4. We implemented a self-supervised clustering module that iteratively refines cluster assignments.

Across 18 diverse scRNA-seq datasets, scAURA consistently achieved strong clustering performance by outperforming most state-of-the-art methods and obtaining the best overall rankings in both ARI and NMI. It also showed robust behavior under increasing dropout levels and maintained stable performance even with high sparsity. In an external Alzheimer’s disease dataset, scAURA not only generated meaningful cell clusters but also identified new marker genes and potential regulators, demonstrating its value for biological discovery. The source code and datasets are available at https://github.com/bozdaglab/scAURA.

## 2 Materials & Methods

### 2.1 Datasets & Data Preprocessing

We collected 18 publicly available scRNA-seq datasets generated across six sequencing platforms (i.e., SMARTer, Smart-seq2, inDrop, Drop-seq, CEL-seq2, and 10x Genomics) and encompassing a range of human and mouse organs, including brain, pancreas, trachea, kidney, liver, and heart. The dataset names are Pollen (Pollen et al., 2014), Camp-Brain (Camp et al., 2015), Camp-Liver (Camp et al., 2017), QS Diaphragm, QS Limb Muscle, QS Trachea, QS Lung, Qx Bladder, Qx Limb Muscle, QS Heart, Qx Spleen (Tabula Muris Consortium et al., 2018), Muraro (Muraro et al., 2016), Klein (Klein et al., 2015), Romanov (Romanov et al., 2017), Adam (Adam et al., 2017), Young (Young et al., 2018), Plasschaert (Plasschaert et al., 2018), and Chen (Chen et al., 2017). The number of profiled cells for each dataset ranges from 301 to 12,089. Detailed characteristics of each dataset, including the number of cells, genes, and cell types, as well as the corresponding sequencing platforms, are summarized in Supplementary Table 1. scRNA-seq data were processed using the Scanpy framework. Gene expression counts were normalized to a total of 10,000 transcripts per cell to correct for differences in sequencing depth. Then log transformation using (1 + *x*) was performed. Highly variable genes (HVGs) were identified using Seurat v3, and the top 500 HVGs were selected for downstream analyses.

### 2.2 Model Architecture

We model single-cell gene expression data on a similarity graph where each node represents a cell and the edges represent cell-cell similarity based on their expression profile. We learn node embeddings using a multi-layer Graph Convolutional Network (GCN) encoder trained via debiased graph contrastive learning. The encoder is regularized with alignment and uniformity objectives to improve representation quality. The learned embeddings are then clustered with *k* -means and refined using a self-supervised clustering technique. The model architecture is shown in Fig. 1. In what follows, we describe each component of scAURA.

**Figure 1.**
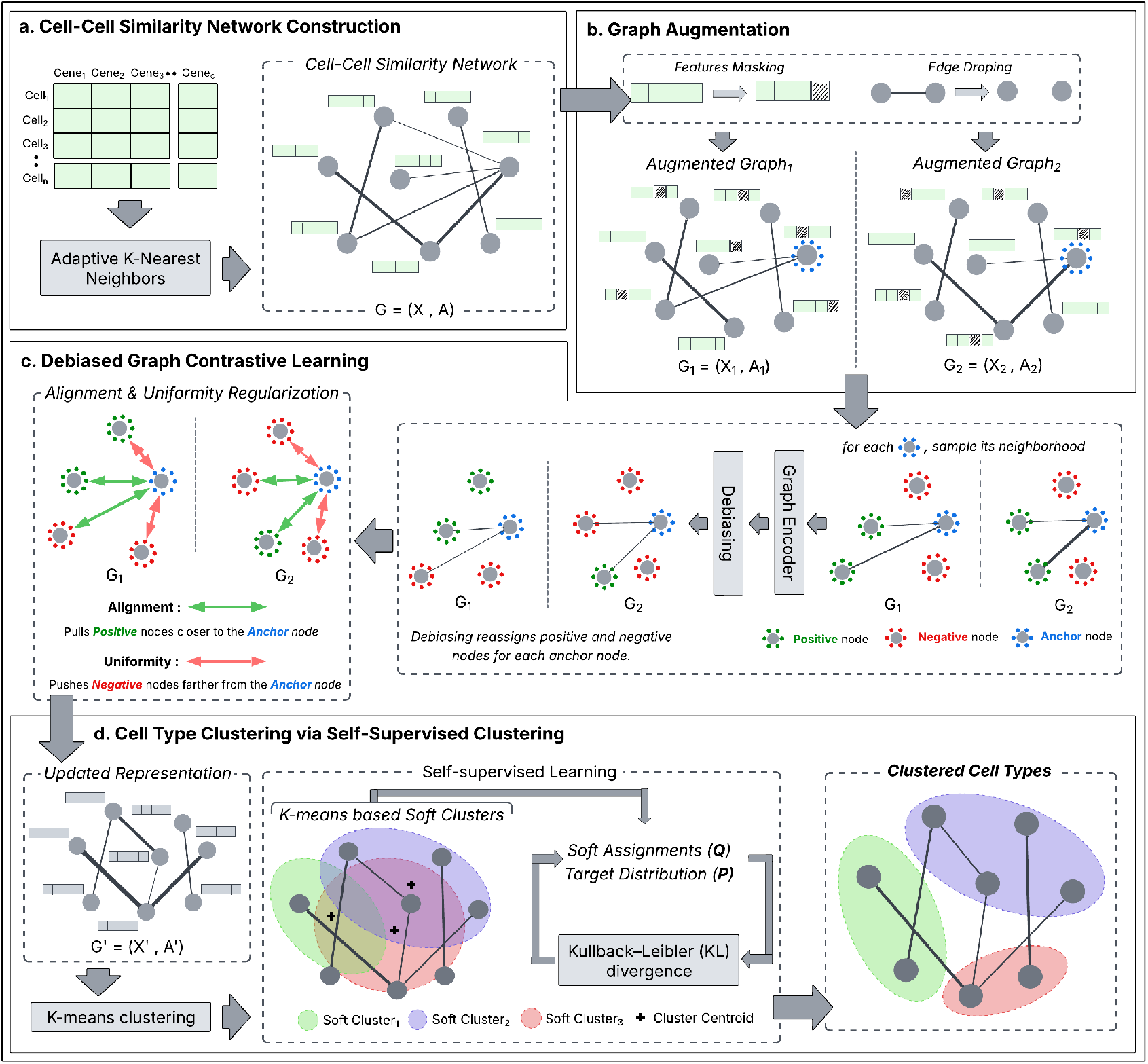
Overview of the scAURA model architecture. **a**. Building cell-cell similarity network. **b**. Augmenting graph with feature masking and edge removing. **c**. Training graph with debiased contrastive learning and regularized by alignment and uniformity. **d**. Self-supervised clustering.

#### 2.2.1 Graph construction from the gene expression matrix

To construct a cell-cell similarity graph, we utilized an adaptive kNN method. Given the gene expression matrix *X* ∈ ℝ^*N ×C*^ , where *N* denotes the number of cells and *C* represents the number of genes, we first apply principal component analysis (PCA) to reduce feature dimensionality. For each cell *i*, we compute the pairwise distances *d*_*ij*_ between cell *i* and all other cell *j* using NearestNeighbors function from scikit-learn library. The top *K*_max_ nearest neighbors are then identified, and an adaptive neighborhood size *k*_*i*_ ≤ *K*_max_ is selected for each cell *i* based on a threshold derived from its sorted neighbor-distance profile.

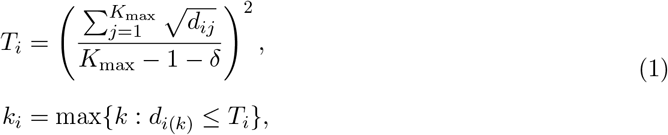

Here, *δ >* 0 controls the adaptivity whee smaller *δ* means large neighborhoods and dense graph and larger *δ* means small neighborhoods and sparse graph. To mitigate spurious edges, we weight edges using the shared nearest-neighbor (SNN) Jaccard similarity. For a directed edge (*i, j*) among the *k*_*i*_ nearest neighbors of cell *i*, the edge weight is defined as

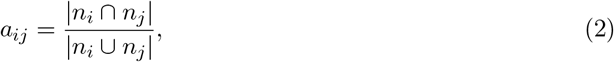

where *n*_*i*_ and *n*_*j*_ denote the sets of the *k*_*i*_ and *k*_*j*_ nearest neighbors of cells *i* and *j*, respectively. This weighting scheme strengthens connections between cells that share many common neighbors while reducing the influence of noisy or isolated relationships.

The weighted adjacency matrix **A** = [*a*_*ij*_] is then constructed directly from the SNN similarity scores *a*_*ij*_. We define the graph *G* = (*V, A, X*) that is used to model cell–cell relationships for downstream representation learning. In *G*, each cell corresponds to a node in *V* , with node features given by the rows of *X*.

#### 2.2.2 Graph encoder

The graph encoder is an *L*-layer Graph Convolutional Network (GCN) with embedding dimension *H* at each layer and nonlinear activation function *σ*. Given a graph *G* = (*V*, **A, X**), the layer-wise propagation is defined as

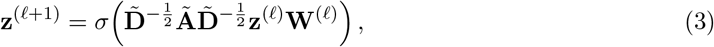

where **z**^(*ℓ*)^ ∈ ℝ^*N ×H*^ denotes node embeddings at layer *ℓ*, **W**^(*ℓ*)^ ∈ ℝ^*H×H*^ is a trainable weight matrix, and **Ã** = **A** + **I**. Here, 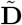 is the corresponding diagonal degree matrix with 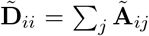. The initial node representations are given by **z**^(0)^ = **X**, and **z**^(*L*)^ serves as the final learned representation after *L* graph convolutional layers.

#### 2.2.3 Projection head

A two-layer projection head maps the final GCN embeddings to a latent space for contrastive learning:

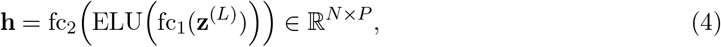

where fc_1_ and fc_2_ denote fully connected layers, and ELU is the Exponential Linear Unit activation function. Here, **h**_*i*_ ∈ ℝ^*P*^ represents the projected embedding of cell *i*, and *P* denotes the dimensionality of the projection space.

#### 2.2.4 Debiased graph contrastive learning

Two graph views are generated through stochastic augmentations, namely edge removing and feature masking where some random edges are dropped and some random node features are masked, respectively based on a probability distribution.

Given two graph views, the shared encoder produces **h**^1^ and **h**^2^. Positive pairs correspond to the same cell across views, while all other nodes serve as negatives. We optimize the debiased InfoNCE objective with temperature *τ* and debiasing factor *τ* ^+^:

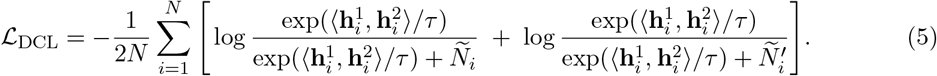

where the debiased negative partitions are

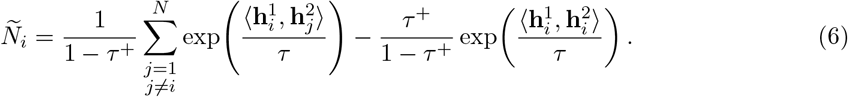

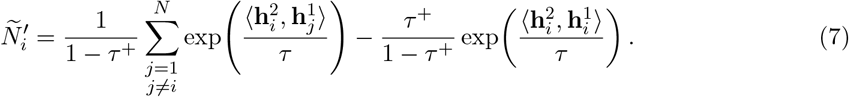

Here, 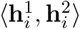 means the dot product between these two vectors, and *τ* ^+^ ∈ [0, 1) is the debiasing term that estimates the probability of a randomly sampled “negative” is actually a false negative, and this debiasing term in Equations (6) and (7) down-weights the corresponding negative contribution accordingly.

#### 2.2.5 Loss modification with alignment and uniformity

To strengthen the contrastive objective, we add two extra losses that help the model learn stable and well-structured representations. The *alignment loss* makes the embeddings of the same cell in the two augmented graph views similar, reducing differences caused by augmentation: 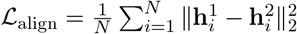. The *uniformity loss* spreads the embeddings out in the representation space, preventing overly compact representations and reducing excessive similarity across all cells: 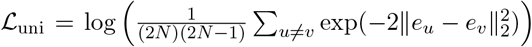 , where *e* denotes the concatenated normalized embeddings of both views. The overall loss function is:

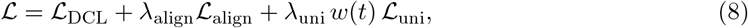

where *w*(*t*) ∈ [0, 1] is a warm-up schedule that gradually increases the weight of the uniformity term during early epochs to prevent over-dispersion. The encoder is then optimized with Adam or stochastic gradient descent (SGD).

#### 2.2.6 Optimizing clustering with self-supervised clustering

After contrastive training, we compute embeddings *z* and perform K-means clustering and determine the optimal number of clusters *K* using the elbow method. The K-means cluster labels are used to obtain initial labels and centroids. Cluster assignments are refined jointly with the encoder using a Student-*t* kernel-based soft assignment:

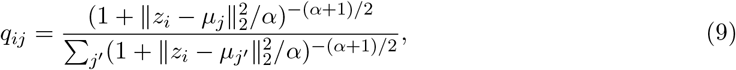

and the target distribution:

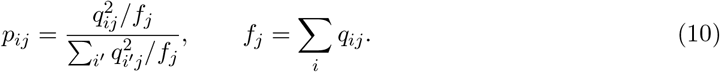

The objective is to minimize the KL divergence:

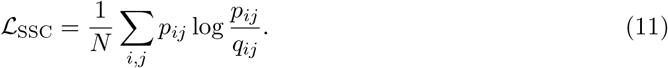

The encoder parameters and cluster centers {*µ*_*j*_} are optimized using SGD. During training, a target distribution is periodically recomputed after a fixed number of iterations to refine the soft cluster assignments and guide the optimization process. The fraction of label changes between consecutive updates are calculated and training is terminated once this fraction falls below a predefined tolerance threshold. Final cluster assignments are evaluated using Adjusted Rand Index (ARI) and Normalized Mutual Information (NMI). The hyperparameters of this model are described in Supplementary Note 1 & Supplementary Table 2.

## 3 Results

### 3.1 scAURA Outperformed State-of-the-Art (SOTA) Methods

We conducted an extensive evaluation of scAURA and compared it to 13 state-of-the-art (SOTA) single-cell clustering algorithms, namely Seurat, SHARP, DCA, DEC, DESC, scziDesk, scCAEs, scVI, scMGCA, scDeepCluster, scGAE, GraphSCC, and scGNN, across 18 benchmark datasets covering six sequencing platforms (see Section 2.1). Clustering performance was assessed using ARI and NMI. To ensure consistency, we ran scAURA on each dataset ten times and computed mean and standard deviation of ARI and NMI. For SOTA methods, the results were obtained from (Yu et al., 2023).

scAURA demonstrated consistent improvements across datasets of various platforms by out-performing the SOTA methods in nine and eight datasets in ARI and NMI, respectively (Fig. 2a, Supplementary Fig. 1, Supplementary Table 3). We computed the average rank of each tool across all 18 benchmark datasets. scAURA achieved the best average rank in both ARI (2.28) and NMI (2.39) (Fig. 2b, Supplementary Table 3). For instance, in the Pollen and QS Diaphragm datasets, which contain 301 and 870 cells, respectively, scAURA achieved performance gains of over 1.5% in ARI compared to the second-best method. Likewise, in Muraro dataset, scAURA improved clustering quality by 2.46%, and 3.26% in ARI and NMI, respectively. These results highlight the model’s capacity to effectively capture complex biological variation and maintain robust clustering performance even when the platform changes. scAURA demonstrated the highest improvements in ARI and NMI on the Camp-Brain, Camp-Liver, Young, and Klein datasets, where the best clustering performance was relatively poor. Specifically, on the Camp-Brain dataset (777 cells) and the Camp-Liver dataset (870 cells), scAURA achieved substantial performance improvements, with gains of 31.4% ARI and 4.4% NMI on Camp-Brain, and 13.6% ARI and 6.8% NMI on Camp-Liver, respectively. Similarly, in Klein, where the best ARI and NMI were only 0.86, our framework improved clustering performance to 0.91 ARI and 0.87 NMI, corresponding to 5.5% ARI and 0.4% NMI improvements. For the datasets where existing methods achieve strong clustering performance (e.g., QS Diaphragm and Qx Bladder), scAURA consistently provided improvements of approximately 1.5%, and 1.2% in ARI and NMI, respectively or very close to the best-performing tool. For example, in Qx Bladder, scAURA achieved an ARI of 0.98 while the best tool achieved 0.99 and matched the NMI performance with the best tool by achieving 0.97. These results indicate that our tool continues to refine clusters even when clustering performance is near saturation. scAURA did not achieve the top performance on a few datasets, likely due to the heterogeneity introduced by different platforms and species. Nevertheless, it consistently ranked second or third and achieved the best overall average rank in terms of ARI among the 13 SOTA tools.

**Figure 2.**
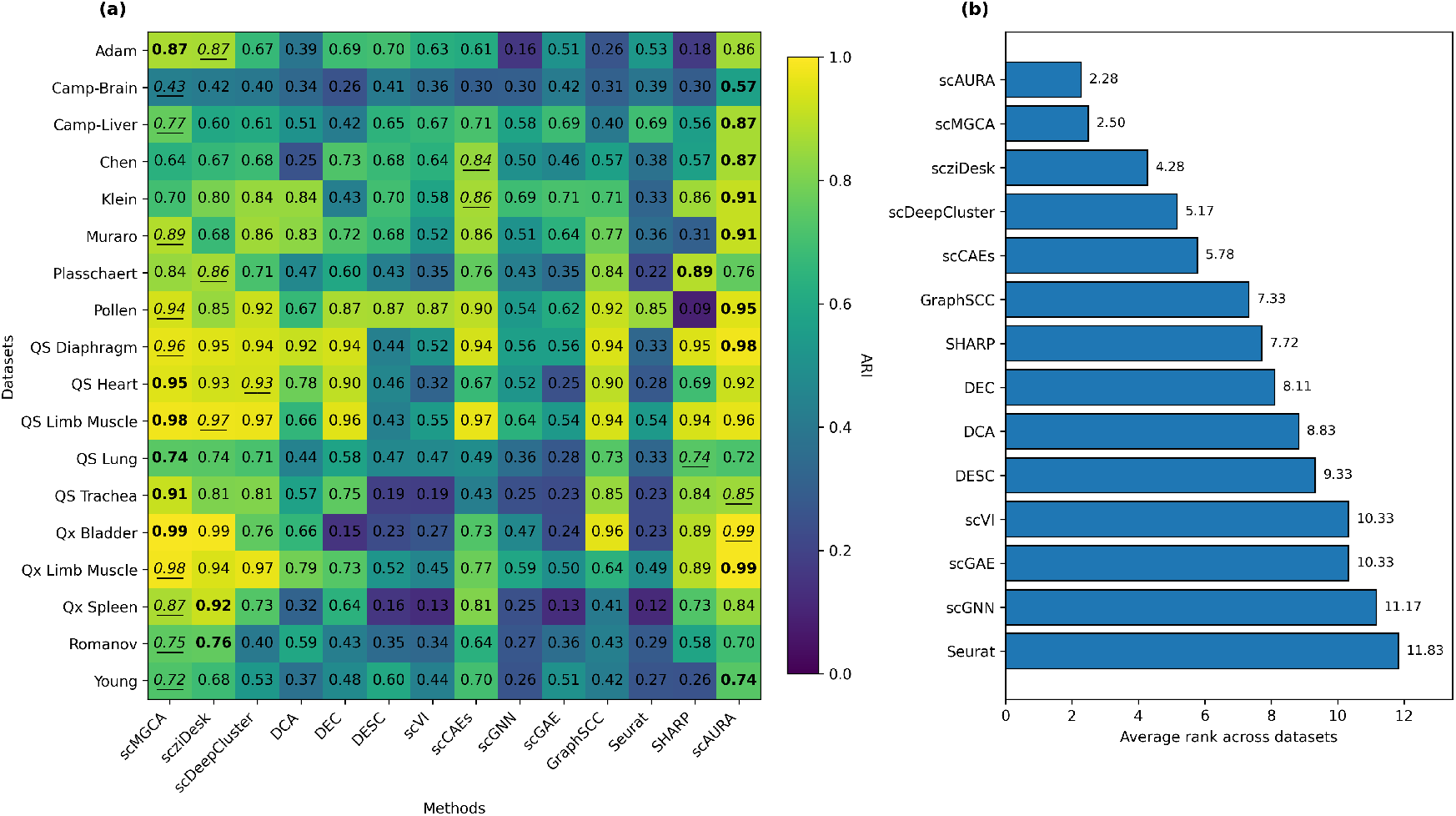
**(a)** The heatmap of Adjusted Rand Index (ARI) scores of SOTA methods and scAURA across 18 benchmark scRNA-seq datasets. For each dataset, the highest and second highest value is shown in **bold** and <u>underlined</u>, respectively. For scAURA, mean ARI across ten runs are shown. **(b)** Average ranking of each tool across datasets based on ARI score (lower is better).

### 3.2 scAURA Demonstrated Robust Clustering Performance in Dropout Correction

Dropout events are a common source of noise in scRNA-seq data, often leading to artificially sparse expression matrices and affecting downstream analyses such as clustering. To evaluate how robust our method is to such dropout-induced sparsity, we compared its clustering performance with eleven deep learning–based clustering approaches under varying levels of simulated dropout. Specifically, we used six representative single-cell datasets obtained from different sequencing platforms and systematically introduced increasing dropout rates to assess how each method responds to missing gene expression information.

Following the simulation protocol described in the DeepImpute study (Arisdakessian et al., 2019), we generated dropout by randomly masking nonzero gene expression values with zeros. The masking probability was gradually increased to emulate dropout levels of 0%, 5%, 10%, 20%, and 50%. This experimental design enabled a controlled evaluation of clustering robustness as data sparsity increased. Across increasing dropout rates (0%–50%), scAURA demonstrated remarkable robustness in both ARI and NMI, showing minimal degradation compared to competing methods (Supplementary Fig. 2 and 3). In the QS Diaphragm dataset, the ARI and NMI values of scMGCA dropped from 0.96 to 0.93 (3.1%) and from 0.94 to 0.89 (5.3%), respectively, as dropout increased to 50%. In contrast, scAURA maintained stable performance with ARI but NMI increased from 0.95 to 0.97. Similarly, in the Muraro dataset, while scMGCA’s ARI and NMI decreased from 0.89 to 0.82 (7.9%) and from 0.84 to 0.80 (4.7% decrease), respectively, scAURA sustained strong clustering stability with ARI above 0.91 and NMI above 0.86 across all dropout levels except for 10% dropout rate. These consistent results across datasets confirm that our graph-debiased contrastive learning approach effectively mitigates the adverse effects of dropout by maintaining stable graph connectivity and emphasizing reliable cell-cell relationships.

### 3.3 Ablation Study

We present an ablation study to evaluate the contribution of each module in our framework. Specifically, we compare graph construction using regular KNN versus adaptive KNN, assess performance with and without the Self-Supervised Clustering (SSC) module, and examine the impact of incorporating alignment and uniformity terms in the loss function.

To assess the effect of graph construction on clustering performance, we compared the conventional fixed K-nearest neighbor (KNN) graphs (*K* = 10, 20, 30, 40) to adaptive KNN strategy across three datasets. We observed that the clustering performance under fixed KNN configurations varies considerably with the choice of *K* (Table 1 & Supplementary Table 4), reflecting the sensitivity of traditional graph construction to suboptimal neighborhood size. In contrast, our adaptive KNN approach consistently produced higher performance across all six datasets by dynamically selecting neighborhood sizes based on local graph structure, minimizing the inclusion of spurious edges, and enhancing graph connectivity. We observed that incorporating SSC consistently improved both ARI and NMI across all datasets except ARI in QS Diaphragm dataset where the values remained the same (Table 1 & Supplementary Table 4). The largest improvement for ARI was observed in Adam (32.2%) and NMI in Klein (19.2%), indicating that SSC enhances the model’s ability to capture complex transcriptional variability. We also observed that excluding either or both uniformity and alignment objectives led to a decline in performance (except for the Muraro dataset where removing alignment or both alignment and uniformity improved the result (Table 1 & Supplementary Table 4). These results confirm that SSC, alignment, and uniformity play complementary roles in learning robust and discriminative representations.

**Table 1:** Performance comparison (ARI) of scAURA variants across datasets. Highest and second-highest values in each row are shown in **bold** and <u>underlined</u>, respectively. For the w/KNN column, the values indicate the range of performance for various choices of *K*. For the other columns, mean and standard deviation across ten runs are shown. SSC: Self Supervised Clustering, A: Alignment, U: Uniformity.

| Dataset | w/ KNN | w/o SSC | w/o A | w/o U | w/o (A+U) | scAURA |
| --- | --- | --- | --- | --- | --- | --- |
| Camp-Liver | <u>0.8443–0.8638</u> | 0.7865±0.0552 | 0.7575±0.0420 | 0.7680±0.0950 | 0.7880±0.0358 | **0.8696±0.0596** |
| QS Diaphragm | 0.9381–0.9664 | **0.9783±0.0040** | <u>0.9778±0.0046</u> | 0.9777±0.0045 | 0.9779±0.0046 | **0.9783±0.0040** |
| Muraro | 0.8506–0.8840 | 0.9068±0.0143 | **0.9158±0.0129** | 0.6613±0.0757 | <u>0.9145±0.0133</u> | 0.9124±0.0105 |
| Klein | 0.4812–0.6766 | <u>0.7321±0.0205</u> | 0.7230±0.0033 | 0.7266±0.0014 | 0.7265±0.0017 | **0.9115±0.0123** |
| Adam | 0.5425–0.6850 | 0.6552±0.0743 | 0.6363±0.0885 | <u>0.7237±0.0690</u> | 0.6868±0.0671 | **0.8659±0.0194** |
| Qx Limb Muscle | 0.8575–0.9540 | 0.9523±0.0723 | 0.8220±0.0764 | <u>0.9618±0.0570</u> | 0.9818±0.0033 | **0.9857±0.0000** |

### 3.4 scAURA clusters cell types and infers potential regulators for cell types in Alzheimer’s Disease data

To further demonstrate the utility of scAURA, we applied it to an external single-nucleus RNA-seq dataset from Alzheimer’s disease (AD) patients (GEO accession GSE138852), which includes 13,214 nuclei from six AD and six control brains. scAURA identified six clusters from six ground truth major cell types: microglia (mg), neurons, oligodendrocyte progenitor cells (OPCs), astrocytes, oligodendrocytes (Oligo), and endothelial cells (endo). Differential expression analysis was performed using the Wilcoxon rank-sum test, where each cluster was compared against all remaining cells (e.g., cluster 1 vs. the rest). Genes were ranked by Wilcoxon test scores, reflecting how consistently a gene’s expression is higher in one cluster compared to others, and the top 25 genes per cluster were selected. As shown in Fig. 3a, these top DE genes effectively distinguished five major cell types (i.e., mg, neurons, OPCs, oligo, and astrocytes). DE genes could not effectively identify endo cell type due to its extreme class imbalance, with only 98 cells out of 11,884 (0.8%). The expression distribution of the top DE gene for each cluster (Fig. 3b) revealed clear cell-type specificity—*TNR* in OPCs, *TMEM144* in oligodendrocytes, *PTPRC* in microglia, and *GABRB2* in neurons while *HSPA1A* showed no strong cell-type preference.

**Figure 3.**
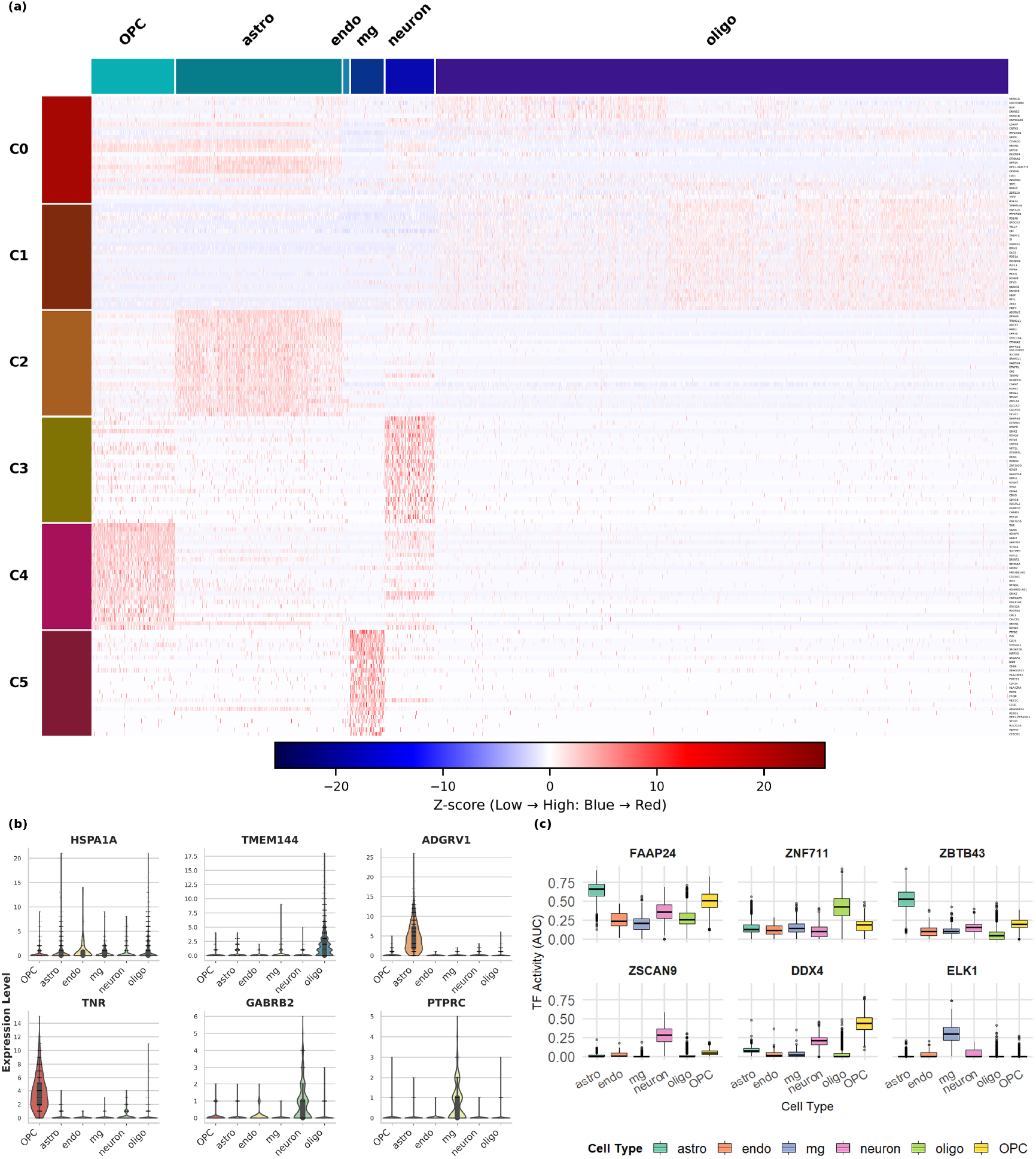
Alzheimer’s disease dataset analysis based on scAURA. **(a)** Heatmap of top 25 Differentially Expressed (DE) genes in each cluster. **(b)** Violin plot of expression levels of top marker genes in each cell cluster. **(c)** Boxplot of AUCell-derived TF activity score of six top regulators across cell types.

To investigate the regulatory control of these DE genes, we applied a R library RcisTarget (Sara Aibar, 2018), which identifies transcription factors (TFs) with enriched binding motifs among the DE genes of each cluster (Supplementary Table 4). This analysis highlighted *FAAP24, ZNF711, ZBTB43, ZSCAN9, DDX4*, and *ELK1* as the top TFs for clusters 0, 1, 2, 3, 4, and 5, respectively. Using AUCell (Aibar et al., 2017), we then quantified regulon activity per cell based on TF–target relationships predicted by RcisTarget. As shown in Fig. 3c, the identified TF regulons exhibit clear cluster-specific activity (except for *FAAP24*), suggesting that their target genes may serve as robust markers for the corresponding cell types.

## 4 Conclusion

We proposed a graph-based debiased contrastive learning framework, scAURA, for single-cell clustering that integrates adaptive graph construction and self-supervised clustering optimization. scAURA outperformed 13 SOTA tools in nine and eight of 18 benchmark datasets spanning six sequencing platforms, in ARI and NMI, respectively. scAURA demonstrated strong generalization across diverse tissue types and sequencing technologies while maintaining robustness to dropout noise, with minimal performance degradation even under 50% simulated dropout on most datasets. Ablation studies revealed that adaptive KNN graph construction, alignment, and uniformity regularization for contrastive loss and the self-supervised clustering module were critical for the model’s success. Furthermore, scAURA successfully identified distinct cell types in an external AD dataset, discovered key marker genes, and inferred their potential transcriptional regulators. Overall, our approach effectively addresses challenges in dropout correction, graph quality, and unsupervised learning, providing a reliable and generalizable solution for single-cell transcriptomic clustering. In future work, we aim to extend our approach to integrate multimodal and spatial omics data for a more comprehensive understanding of cellular heterogeneity.

## Conflicts of interest

The authors declare that they have no competing interests.

## Funding

This work was supported by funds from the National Institute of Health under grant MIRA (R35GM133657) to S.B. and Summer Research Assistantship from the BioDiscovery Institute to J.I.M.R.

## Data availability

All datasets used in this study are publicly available (see Section 2.1).

## Author contributions statement

Conceptualization, J.I.M.R., S.B.; Methodology, J.I.M.R., S.B.; Data collection, J.I.M.R.; Running experiments, J.I.M.R., S.E.; Writing—Original Draft, J.I.M.R.; Writing—Review & Editing, S.B., J.I.M.R.; Visualization, J.I.M.R.; Supervision: S.B.

## Notes

https://github.com/bozdaglab/scAURA

### Competing Interest Statement

The authors have declared no competing interest.

