## supplementary files for "scAURA: Alignment- and Uniformity-based Graph Debiased Contrastive Representation Architecture for Self-Supervised Clustering of Single-Cell Transcriptomics"

Supplementary Note 1: Hyperparameter Description

The training procedure of scAURA can be divided into three stages: (i) graph construction, (ii) representation learning via graph-debiased contrastive learning, and (iii) self-supervised clustering technique. Each stage has a distinct set of hyperparameters that control model behavior and optimization. Graph construction hyperparameters define how cell–cell relationships are established from gene expression data. Contrastive learning hyperparameters regulate the architecture of the graph encoder, graph augmentations, and the modified debiased contrastive objective used to learn robust latent representations. Self-supervised clustering hyperparameters control the refinement of cluster assignments through iterative optimization. Supplementary Table 2 summarizes all hyperparameters, their roles, and the values used across the 18 datasets.

Supplementary Table 1: Description of 18 real scRNA-seq datasets used in this study.

| **Dataset** | **Organ** | **# Cells** | **# Genes** | **# Cell Types** | **Platform** |
| --- | --- | --- | --- | --- | --- |
| Pollen | Tissues | 301 | 21,721 | 11 | SMARTer |
| Camp-Brain | Brain | 734 | 18,927 | 6 | SMARTer |
| Camp-Liver | Liver | 777 | 19,020 | 7 | SMARTer |
| QS Diaphragm | Diaphragm | 870 | 23,341 | 5 | Smart-seq2 |
| QS Limb Muscle | Limb Muscle | 1,090 | 23,341 | 6 | Smart-seq2 |
| QS Trachea | Trachea | 1,350 | 23,341 | 2 | Smart-seq2 |
| QS Lung | Lung | 1,676 | 23,341 | 11 | Smart-seq2 |
| Muraro | Pancreas | 2,122 | 19,046 | 9 | CEL-seq2 |
| Qx Bladder | Bladder | 2,500 | 23,341 | 4 | 10x Genomics |
| Klein | Embryonic Stem Cell | 2,717 | 24,047 | 4 | inDrop |
| Romanov | Hypothalamus | 2,881 | 21,143 | 7 | SMARTer |
| Adam | Kidney | 3,660 | 23,797 | 8 | Drop-seq |
| Qx Limb Muscle | Limb Muscle | 3,909 | 23,341 | 6 | 10x Genomics |
| QS Heart | Heart | 4,365 | 23,341 | 8 | Smart-seq2 |
| Young | Kidney | 5,685 | 33,658 | 11 | 10x Genomics |
| Plasschaert | Trachea | 6,977 | 28,205 | 8 | inDrop |
| Qx Spleen | Spleen | 9,522 | 23,341 | 5 | 10x Genomics |
| Chen | Brain | 12,089 | 23,284 | 46 | Drop-seq |

Supplementary Table 2**.** Hyperparameters used in scAURA and their roles.

| **Hyperparameter** | **Description** | **Values** |
| --- | --- | --- |
| **Graph Construction** | | |
| n_pca | Number of principal components retained when PCA is applied. | [10,20,30,40,50,60,70] |
| $K_{max}$ | Maximum neighborhood size for adaptive kNN graph construction | [10,20,30,40,50] |
| $\delta$ | Adaptivity parameter controlling how neighborhood sizes are adjusted per cell based on distance distributions. | [-2, -1, 0.5,1,2] |
| dist_metric | Distance metric used to compute cell-cell similarity during graph construction. | [euclidean, cosine] |
| **Graph Debiased Contrastive Learning** | | |
| hidden_dim | Dimensionality of hidden layers in the GCN encoder | [16,32,64] |
| P | Output dimensionality of the projection head used for contrastive learning. | [8,16,32,64] |
| num_gcn_layers | Number of graph convolutional layers in the encoder. | [2,3] |
| activation | Nonlinear activation function used in the GCN layers | [sigmoid, relu, tanh] |
| edge_dropout_prob | Edge dropout probability used for graph augmentation | [0.1,0.2,0.3,0.4,0.5] |
| feature_masking_prob | Feature masking probability applied to node features | [0.1,0.2,0.3,0.4,0.5] |
| $\tau$ | Temperature parameter in the contrastive loss that controls the sharpness of similarity distributions. | [0.5,0.6,0.7,0.8] |
| $\tau^{+}$ | Debiasing factor that estimates the probability of false negatives in contrastive learning. | [0.1,0.2,0.3,0.4] |
| lr | Learning rate for optimizing model parameters. | [0.01,0.001,0.0001] |
| epochs | Number of training epochs | [50,100,200,300,400] |
| $\lambda_{align}$ | Weight of the alignment loss | [0, 0.1,0.2,0.3,1] |
| $\lambda_{uni}$ | Weight of the uniformity loss. | [0, 0.1,0.2,0.3,1] |
| uniform_warmup_epochs | Number of warm-up epochs before fully applying the uniformity loss to avoid early over-dispersion. | [10,20,30] |
| **Self-Supervised Clustering** | | |
| ssc_batch_size | Mini-batch size used during the self-supervised clustering refinement stage. | [8,16,32,64,128] |
| $\alpha$ | Degrees-of-freedom parameter of the Student-t distribution used for soft cluster assignment. | [1] |
| ssc_lr | Learning rate used during self-supervised clustering refinement stage. | [0.001,0.001] |
| ssc_max_epochs | Maximum number of epochs for clustering refinement. | [300,400,500] |
| ssc_update_iters | Number of iterations between updates of the target distribution in self-supervised clustering. | [10,20,50] |
| tol | Tolerance threshold for convergence based on the fraction of changed cluster assignments. | [0.005,0.0005] |

Supplementary Table 3: Summary of ARI and NMI Percentage changes comparing to best tool

| **Datasets** | **Platforms** | **Comparing Best ARI Method** | **Comparing Best NMI Method** | **scAURA ARI**  **(mean**±std. dev) | **scAURA NMI**  **(mean**±std. dev) | **ARI percent change** | **NMI percent change** |
| --- | --- | --- | --- | --- | --- | --- | --- |
| Pollen | SMARTer | scMGCA | scVI | 0.9512±  0.0020 | 0.9455±  0.0023 | 1.5263 | 0.0106 |
| Camp-Brain | SMARTer | scMGCA | scMGCA | 0.5668± 0.0127 | 0.5834± 0.0128 | 31.3862 | 4.3836 |
| Camp-Liver | SMARTer | scMGCA | scMGCA | 0.8696± 0.0596 | 0.8984± 0.0249 | 13.6583 | 6.8252 |
| QS Diaphragm | Smart-seq2 | scMGCA | scMGCA | 0.9783± 0.0040 | 0.9496± 0.0158 | 1.5466 | 1.172 |
| QS Limb Muscle | Smart-seq2 | scMGCA | scMGCA | 0.9622± 0.0072 | 0.9395± 0.0076 | -1.5048 | -2.0029 |
| QS Trachea | Smart-seq2 | scMGCA | scMGCA | 0.8518± 0.0346 | 0.7779± 0.0471 | -6.8154 | -11.026 |
| QS Lung | Smart-seq2 | scMGCA | scMGCA | 0.7162± 0.0298 | 0.7827± 0.0243 | -3.581 | -3.0956 |
| Muraro | CEL-seq2 | scMGCA | scMGCA | 0.9124± 0.0105 | 0.8690± 0.0125 | 3.0146 | 3.7734 |
| Qx Bladder | 10x | scMGCA | scMGCA | 0.9874± 0.0026 | 0.9703± 0.0067 | -0.3331 | -0.9494 |
| Klein | inDrop | scCAEs | SHARP | 0.9115± 0.0123 | 0.8730± 0.0116 | 5.4855 | 0.4025 |
| Romanov | SMARTer | scziDesk | scMGCA | 0.6951± 0.0385 | 0.6956± 0.0243 | -8.5756 | -5.309 |
| Adam | Drop-seq | scMGCA | scMGCA | 0.8659± 0.0194 | 0.8630+-0.117 | -1.001 | 0.6626 |
| Qx Limb Muscle | 10x | scMGCA | scMGCA | 0.9857± 0.0000 | 0.9681± 0.0000 | 0.4279 | 0.4983 |
| QS Heart | Smart-seq2 | scMGCA | scMGCA | 0.9204± 0.0138 | 0.8690± 0.0141 | -3.1464 | -3.6906 |
| Young | 10x | scMGCA | scMGCA | 0.7417± 0.0344 | 0.8052± 0.0185 | 2.3034 | -0.9107 |
| Plasschaert | inDrop | SHARP | SHARP | 0.7610± 0.0194 | 0.7380± 0.0080 | -14.504 | -8.4708 |
| Qx Spleen | 10x | scziDesk | scziDesk | 0.8432± 0.0220 | 0.7307± 0.0259 | -8.3179 | -11.847 |
| Chen | Drop-seq | scCAEs | scCAEs | 0.8705± 0.0064 | 0.7833± 0.0026 | 3.6062 | -5.7287 |

Supplementary Table 4: Performance comparison (NMI) of scAURA variants across datasets. Highest and second-highest values in each row are shown in bold and underlined, respectively. For the w/ KNN column, the values indicate the range of performance for various choices of K. For the other columns, mean and standard deviation across ten runs are shown. SSC: Self Supervised Clustering, A: Alignment, U: Uniformity

| **Dataset** | **w/ KNN** | **w/o SSC** | **w/o A** | **w/o U** | **w/0 (A+U)** | **scAURA** |
| --- | --- | --- | --- | --- | --- | --- |
| Camp-Liver | 0.8760-0.8844 | 0.8600±  0.0219 | 0.8534±  0.0227 | 0.8559±  0.0402 | 0.8660±  0.0142 | **0.8984**±  **0.0249** |
| Qs Diaphragm | 0.9214-0.9464 | 0.9495±  0.0158 | 0.9487±  0.0155 | 0.9484±  0.0156 | 0.9490±  0.0160 | **0.9496**±  **0.0158** |
| Muraro | 0.8344-0.8511 | 0.8642±  0.0164 | 0.8737±  0.0135 | 0.8040±  0.0426 | **0.8738**±  **0.0159** | 0.8690±  0.0125 |
| Klein | 0.5465-0.6396 | 0.7322±  0.0153 | 0.7250±  0.0041 | 0.7280±  0.0018 | 0.7275±  0.0024 | **0.8730**±  **0.0116** |
| Adam | 0.6663-0.7535 | 0.7613±  0.0399 | 0.7450±  0.0542 | 0.7891±  0.0471 | 0.7692±  0.0498 | **0.8566**±  **0.0131** |
| Qx Limb Muscle | 0.9024-0.9499 | 0.9444±  0.0388 | 0.8859±  0.0360 | 0.9499±  0.0308 | 0.9612±  0.0053 | **0.9681**±  **0.0000** |

Supplementary Table 4: Cluster specific Transcription Factor (TF) obtained from RcisTarget and their target genes from the top 25 DE genes

| cluster | motif | TF | NES | # Gene | Enriched Genes among top25 DE gene |
| --- | --- | --- | --- | --- | --- |
| 0 | hdpi__C19orf40 | FAAP24 | 4.74 | 8 | *ADGRB3, CTNND2, DHCR24,DPP10, GPM6A, LINC00486, PARK2, ZBTB20* |
| 1 | transfac_pro__M05999 | ZNF711 | 5.51 | 9 | *CLDN11, DNAJC6, HHIP, PDE1A, PDE4B, PTPRK, QKI, RNF220, ZEB2* |
| 2 | hdpi__ZBTB43 | ZBTB43 | 5.35 | 10 | *ADCY2, BMPR1B, FGFR3, GABRB1, GPM6A, LSAMP, NHSL1, SLC1A3, SPARCL1, TPD52L1* |
| 3 | hdpi__ZNF193 | ZSCAN9 | 6.06 | 19 | *ADGRL2, CCSER1, CDH18, CDH9, CHRM3, CNTN4, FSTL5, GABRB2, GALNTL6, GRIK2, HCN1, KCNH7, KCNJ3, KCNQ5, MYT1L, ROBO2, SYNPR, ZNF385B, ZNF385D* |
| 4 | hdpi__DDX4 | DDX4 | 5.23 | 16 | *BRINP3, CHL1, CNTNAP5,COL11A1, FGF12, GRID2, GRIK2, GRM7, MIR3681HG, NRXN1, NTNG1, SCN1A, SCN3A, SEMA5A, TNR, VCAN* |
| 5 | transfac_pro__M01204 | ELK1 | 5.94 | 9 | *ARHGAP15, C1QB, C1QC, CX3CR1, INPP5D, MEF2C, PLA2G4A, PTPRC, ST6GAL1* |

Supplementary Fig 1. (a) Heatmap of NMI scores on 18 datasets. (b) Average ranking of SOTA tools across 18 datasets.

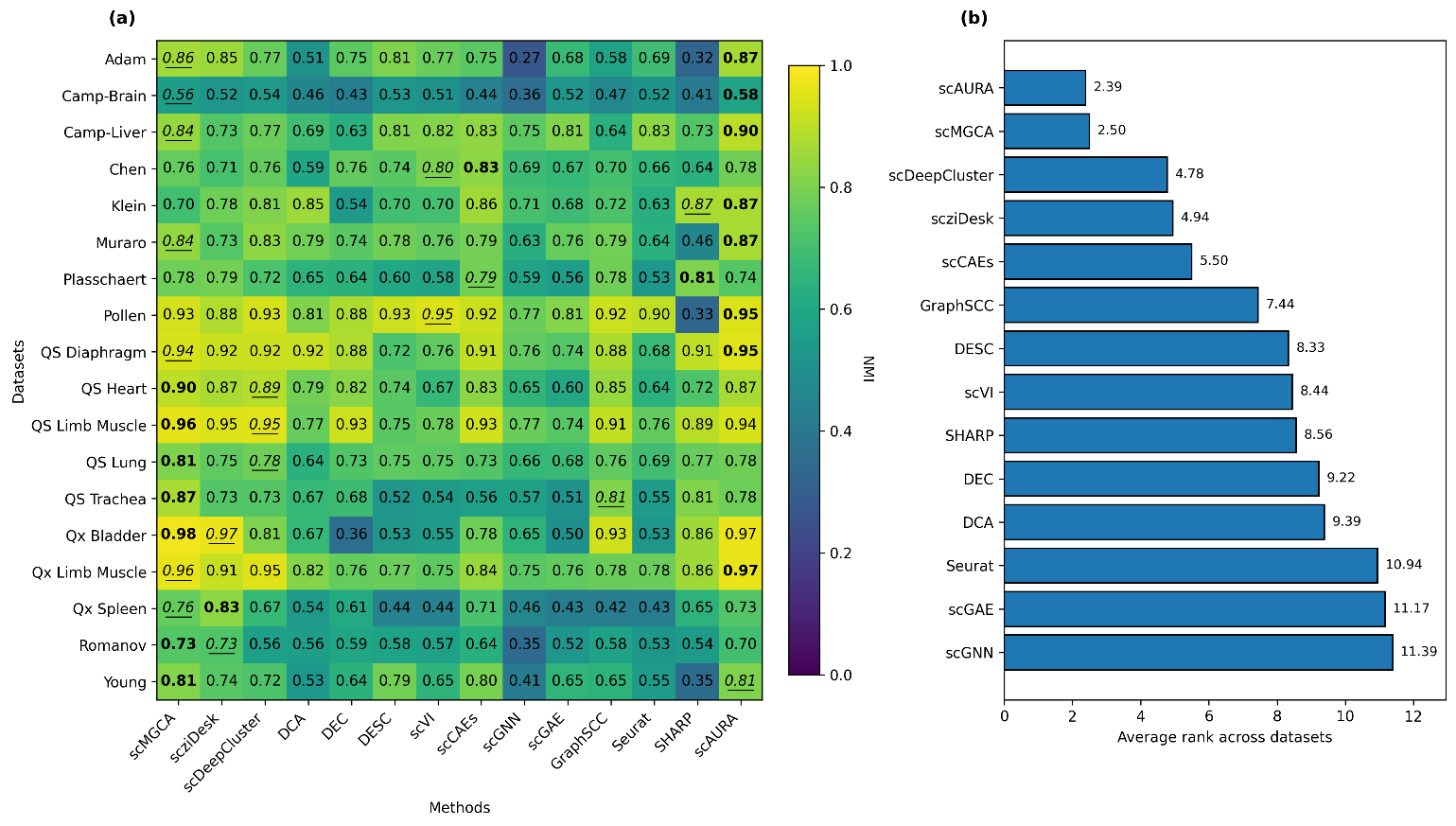

Supplementary Fig. 2. Comparison of ARI under varying simulated dropout rates (0%,5%,10%,20%,50%).

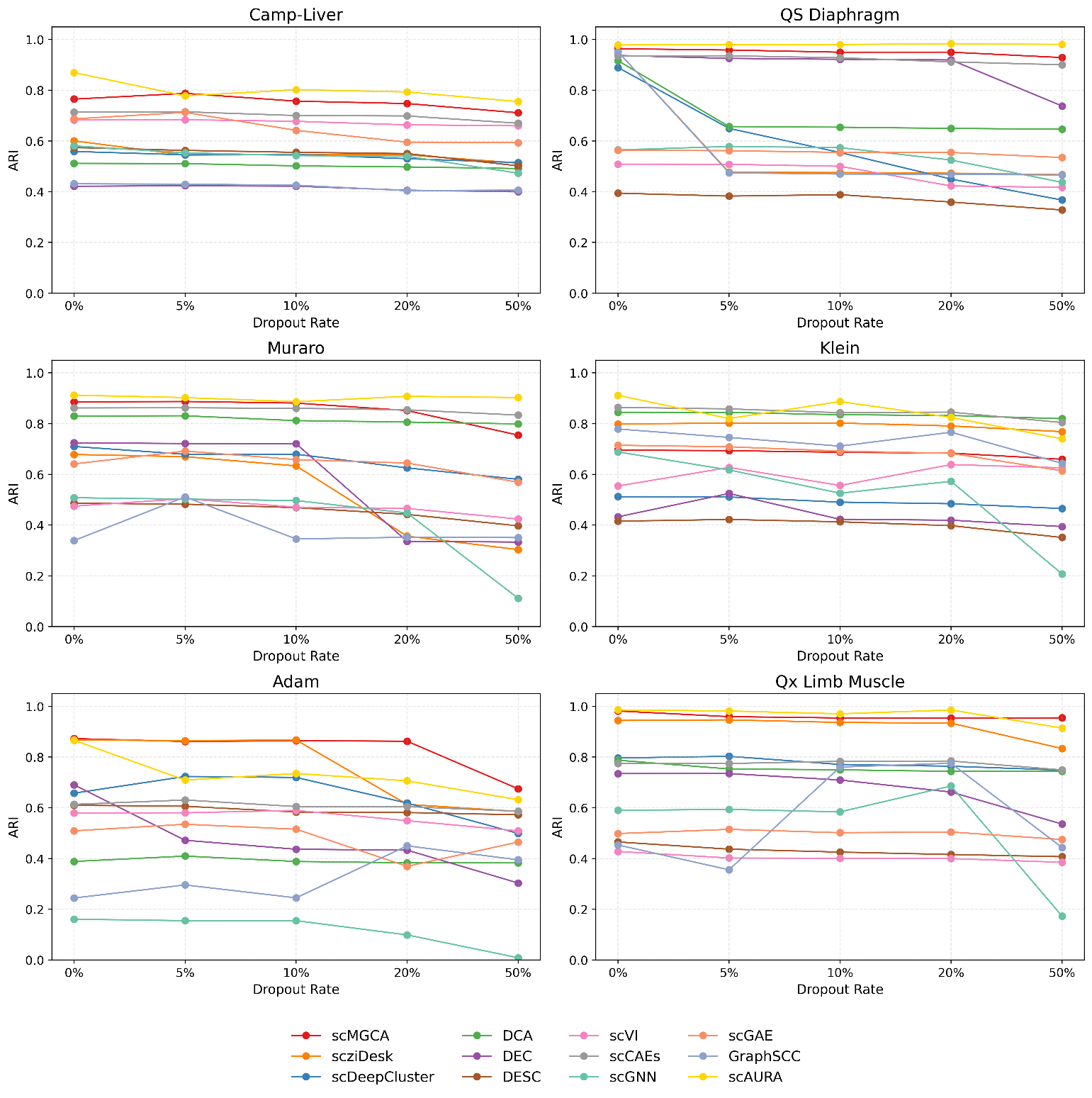

Supplementary Fig 3. Comparison of NMI under varying simulated dropout rates (0%,5%,10%,20%,50%).

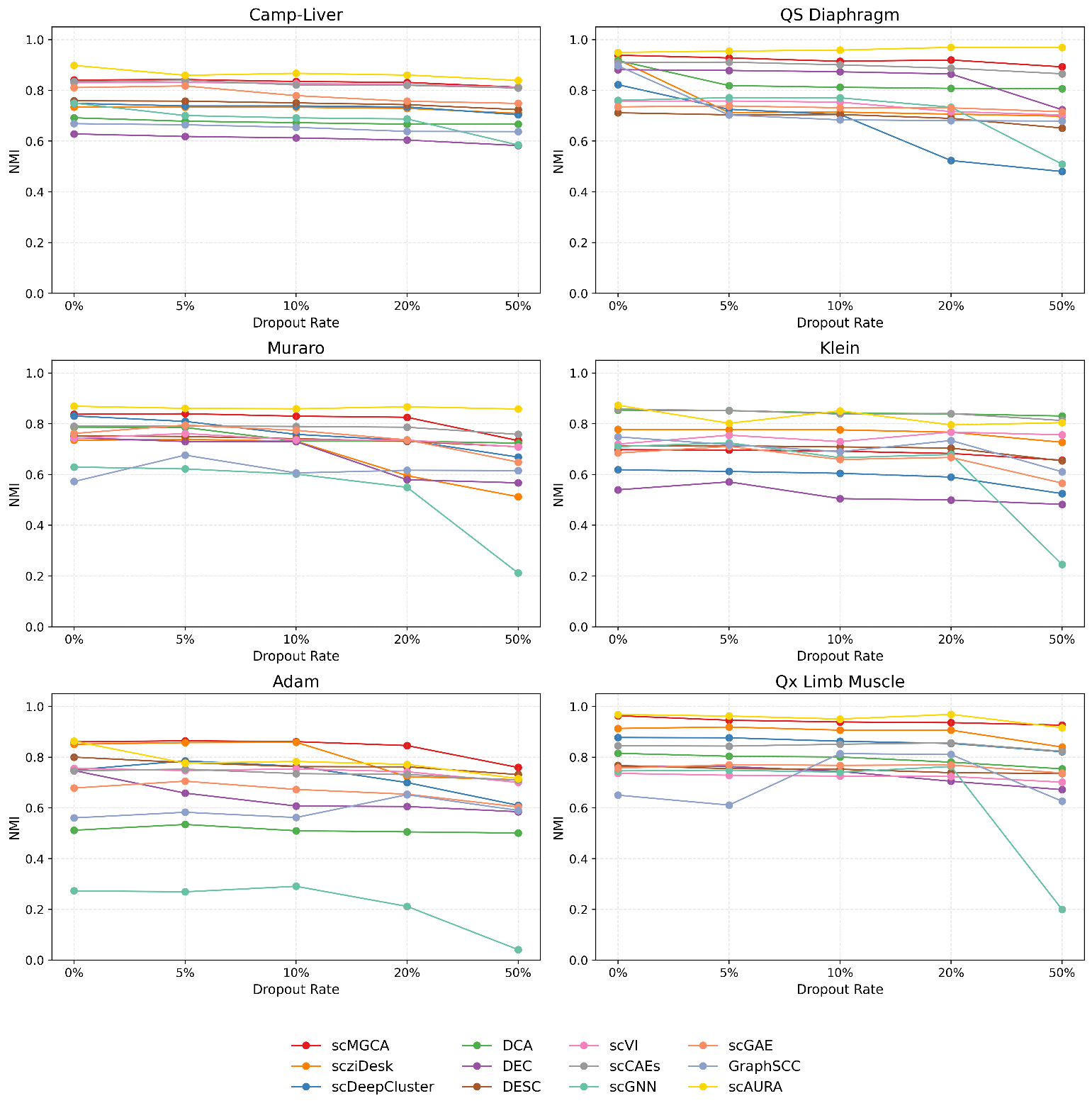

Supplementary Fig 4. Violin Plot of gene expression values for the second most differentially expressed gene in each cluster.

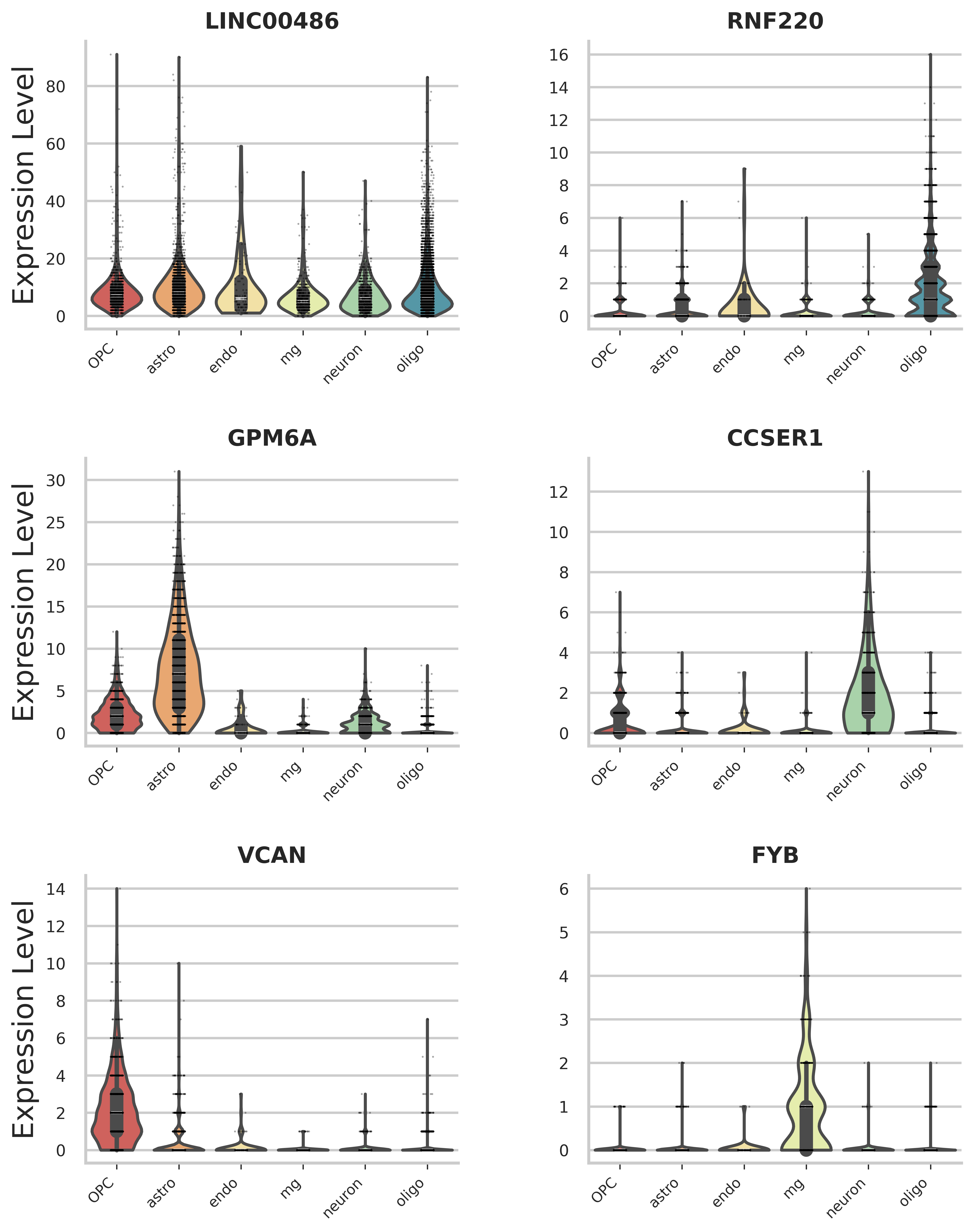
